# Lipid-mediated dimerization of membrane-anchored c-Src is driven by a cluster of lysine residues in the N-terminal SH4 domain

**DOI:** 10.1101/2022.05.31.494233

**Authors:** Irrem-Laareb Mohammad, Javier Carvajal, Alejandro Fernández, Marta Taulés, Elise Fourgous, Yvan Boublik, Anabel-Lise Le Roux, Serge Roche, Miquel Pons

**Affiliations:** Biomolecular NMR laboratory, Department of Inorganic and Organic Chemistry, Universitat de Barcelona (UB), Baldiri Reixac 10-12, 08028 Barcelona, Spain; Centres Científics i Tecnològics (CCiTUB), Universitat de Barcelona (UB), Baldiri Reixac 10-12, 08028 Barcelona, Spain; CNRS UMR5237, University of Montpellier, CRBM, 1919 Rue de Mende, 34000 Montpellier, France; Equipe labellisée Ligue Contre le Cancer, CRBM, University of Montpellier, CNRS, 34000 Montpellier, France; Institut de Bioenginyeria de Catalunya, Baldiri Reixac 15-21, 08028 Barcelona, Spain

## Abstract

The membrane-anchored c-Src tyrosine kinase mediates signaling from a wide range of cell surface receptors controlling cell growth, adhesion, and survival. c-Src deregulation is associated with cancer. Dimerization appears to be a novel layer of regulation through a yet unclear mechanism. Binding of c-Src tyrosine kinase to the plasma membrane is mediated by the myristoylated and strongly positively charged N-terminal SH4 domain. Although activation of c-Src is known to require phosphorylation by a second c-Src molecule, electrostatic repulsion between the charged residues was considered to prevent dimerization. Here we show that a cluster of positively charged lysine residues in c-Src SH4 domain not only does not prevent dimerization but, in fact, enhances it through a lipid-mediated process. Dimerization not only depends on the number of positive charges but also on their position and the nature of the charged residues. Replacement of lysine by arginine increases dimerization in vitro and in vivo and, in HEK293T cells, causes a two-fold increase in tyrosine phosphorylation. Lipid mediated protein-protein interactions induced by clusters of basic residues may represent a general mechanism for modulating cell signaling, consistent with the abundance of positively charged residues in the juxta membrane region of many signaling proteins.

## Introduction

Non-receptor protein kinases of the Src family (SFK) participate in signaling pathways that regulate cell growth, proliferation, differentiation, adhesion and migration (1). Overactivation of c-Src, the prototypical SFK, is associated to poor prognosis in many cancers (2,3). The N-terminal disordered myristoylated SH4 and Unique domains along with the adjacent globular SH3 domain form the Src N-terminal Regulatory Element (SNRE). The key feature of the SNRE is the arrangement of an interdomain fuzzy complex in which the disordered SH4 and Unique domains are condensed around the SH3 domain, while retaining the extensive dynamics typical of intrinsically disordered regions (4,5). In human colon rectal cancer cells, the SNRE has been revealed to have a relevant role in the c-Src mediated oncogenic signaling (6). All SFK members are anchored to membranes through the N-terminal myristoylated SH4 domain. Membrane association is essential for the transforming capacity of the viral form of Src (7). Since the insertion of the single myristoyl group accounts for barely 8 kcal mol^-1^ (8), the majority of SFK, except c-Src, Blk and Hck A, contain at least a second fatty acyl chain (palmitoyl) to ensure stable lipid binding. In c-Src the second membrane binding motif includes positively charged residues in the SH4 domain that interact electrostatically with negatively charged lipids (8–10).

Although it is known that c-Src activation involves trans-phosphorylation by a second c-Src molecule (11), membrane anchored c-Src has been generally assumed to remain monomeric. Nonetheless c-Src dimers have been recently described by us and others (12–15). Membrane interaction is required for c-Src dimerization and results in persistent binding (12), in contrast to the reversible binding observed in c-Src monomers. While the essential role of c-Src SH4 domain as a lipid anchor is unquestionable, recent evidence suggests that the SH4 domain plays other roles in c-Src regulation, by being part of the fuzzy complex involving the Unique and SH3 domains (4,5,16) or by interaction with the kinase domain (17). Here we explore the participation of the SH4 domain in lipid-mediated c-Src dimerization and demonstrate the surprising result that, in spite of the expected electrostatic repulsion, the cluster of positively charged lysine residues in the SH4 domain of c-Src is essential for c-Src dimerization. Replacement of these lysine residues by arginine further enhances dimerization in vitro and in vivo and results in a two-fold increase tyrosine phosphorylation in HEK293T cells. These results provide a new view for the role of positive charges in myristoylated c-Src, beyond the classical model of enhanced monomeric binding by electrostatic interaction with negatively charged lipids.

Although dimers represent a minor fraction of membrane-anchored c-Src, their much slower release and the possibility for enhanced trans-activation, suggest that they could represent enhanced signaling hot spots. These finding also highlight the important role of lipids in the regulation of a non-receptor protein kinase and suggests an important role of lipids, charged or neutral, in mediating protein-protein interactions through the abundant juxtamembrane charged clusters (18).

## Results

### c-Src dimerization is modulated by K5, K7 and K9 in the SH4 domain

In prior studies, we detected and quantified the dimeric form of myristoylated SNRE (aka myrUSH3) on supported lipid bilayers (SLB) through an antibody capture protocol using Surface Plasmon Resonance (SPR) (12). The method exploits the fact that the major monomeric myrUSH3 fraction binds reversibly and therefore can be washed away from the membrane, thus enabling the selectively detection of the minor form that remains permanently anchored to the immobilized lipid bilayer.

The complete protocol (Fig. 1A) consists of: 1) liposome capture, 2) myristoylated protein binding (including reversible and irreversible bound forms) 3) washing of the major monomeric fraction, 4) antibody binding to the remaining, minor persistently bound fraction and 5) regeneration with detergents to strip off the captured liposomes. Persistently bound c-Src dimers represent a minor fraction of the steady-state bound protein, thus the response is amplified with the antibody capture, so that the relative amount of dimer formed by c-Src variants can be accurately quantified. Fig. 1B shows a representative example using myrUSH3 variants detected with an antiHisTag antibody.

**Figure 1.**
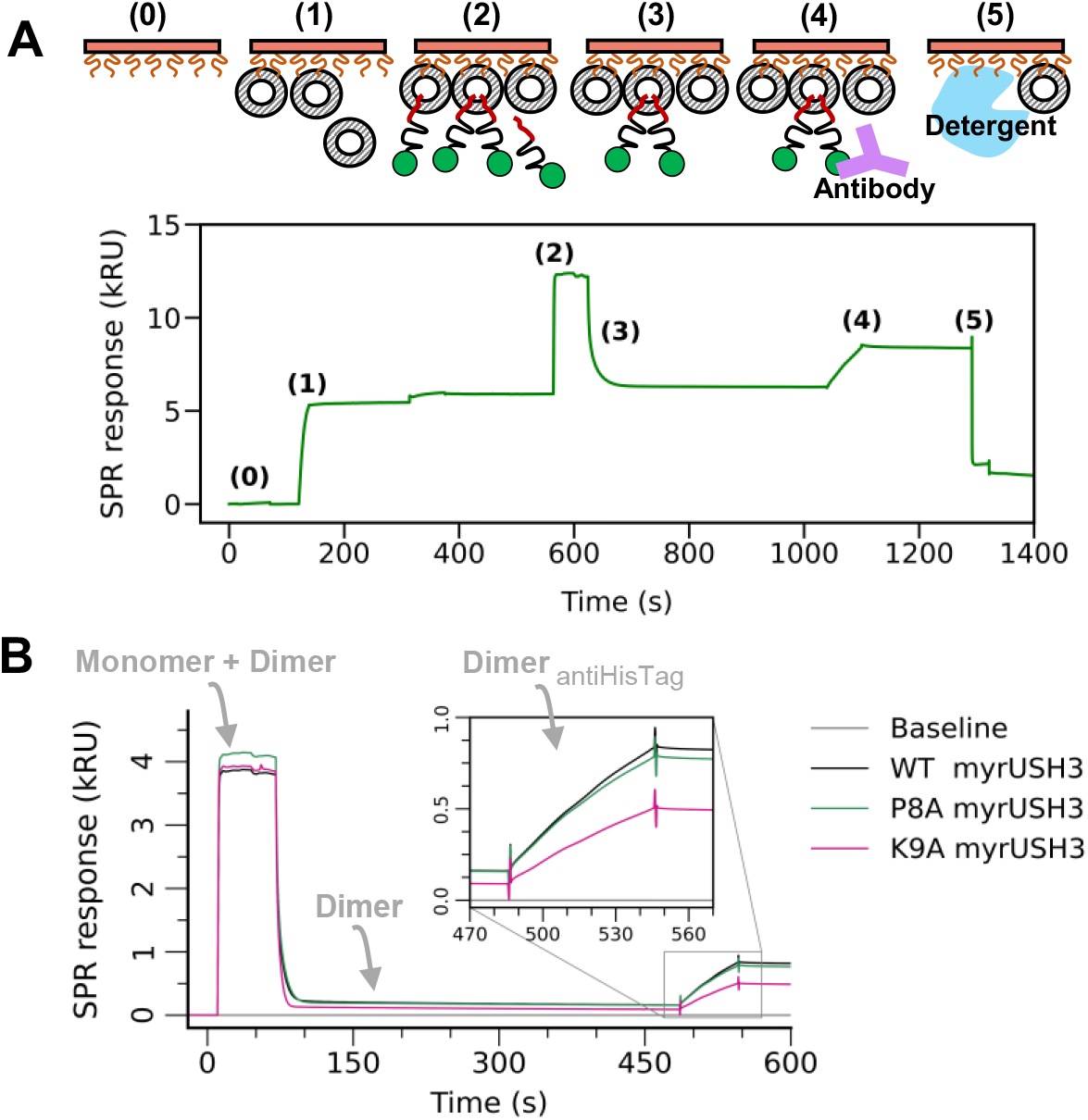
(A) Overview of the protocol to detect the membrane bound minor c-Src self-associated fraction using SPR. (B) Representative SPR sensorgrams displaying the effect of mutations P8A and K9A on the dimerization of myrUSH3 on negatively charged liposomes DOPC:DOPG (2:1). An antiHisTag is used to detect the minor self-associated myrUSH3 fraction.

Dwivedi et al. (19) using short myristoylated peptides showed that residues 2-9 of the SH4 domain are enough to enable oligomerization on the membrane. Thus, we decided to perform an alanine-scan by SPR of residues 3 to 9 in the context of the 150 residues long SNRE to identify the sequence determinants of c-Src dimerization when bound to zwitterionic or negatively charged supported lipid bilayers. The zwitterionic lipid was 1,2-dioleoyl-sn-glycero-3-phospho choline (DOPC) and the negative charged bilayers were formed by a 2:1 mixture of DOPC and 1,2-dioleoyl-sn-glycero-3-phospho (1’-rac glycerol) (DOPG). The two types of membranes are in the liquid disordered phase and the DOPC:DOPG mixture do not form separate phases in the conditions of the experiment (20). We used an antibody directed to the His_6_Tag located at the C-terminus to minimize the interference with the membrane-anchoring region. The primary response is proportional to the amount of membrane-anchored c-Src in steady-state conditions in the presence of a constant concentration of soluble protein. The response to the antibody is proportional to the dimer population. The two SPR responses are not directly comparable, thus, only the relative responses of each of the mutants can be compared.

Fig. 2A and 2B show the relative SPR responses of the individual mutants with respect to wild type (WT) myrUSH3 in zwitterionic and negatively charged immobilized liposomes respectively. Green bars represent the relative amount of monomeric and dimeric myrUSH3 while the orange bars reflect the relative dimer concentration. The absolute concentration of dimer is only a few percent of the total immobilized protein (Fig. 1B). Replacing lysine residues in positions 5 and 7 by alanine reduces dimer formation by around 50% on negatively charged lipid bilayers and by 30-40% in neutral lipids. A similar decrease in dimer formation is observed in the K9A and S3A mutants bound to negatively charged lipids. Interestingly, the residues whose mutation cause the maximum effect on dimerization are located on alternating positions. This includes the three alternating lysine residues at positions 5, 7 and 9 but also serine 3.

**Figure 2.**
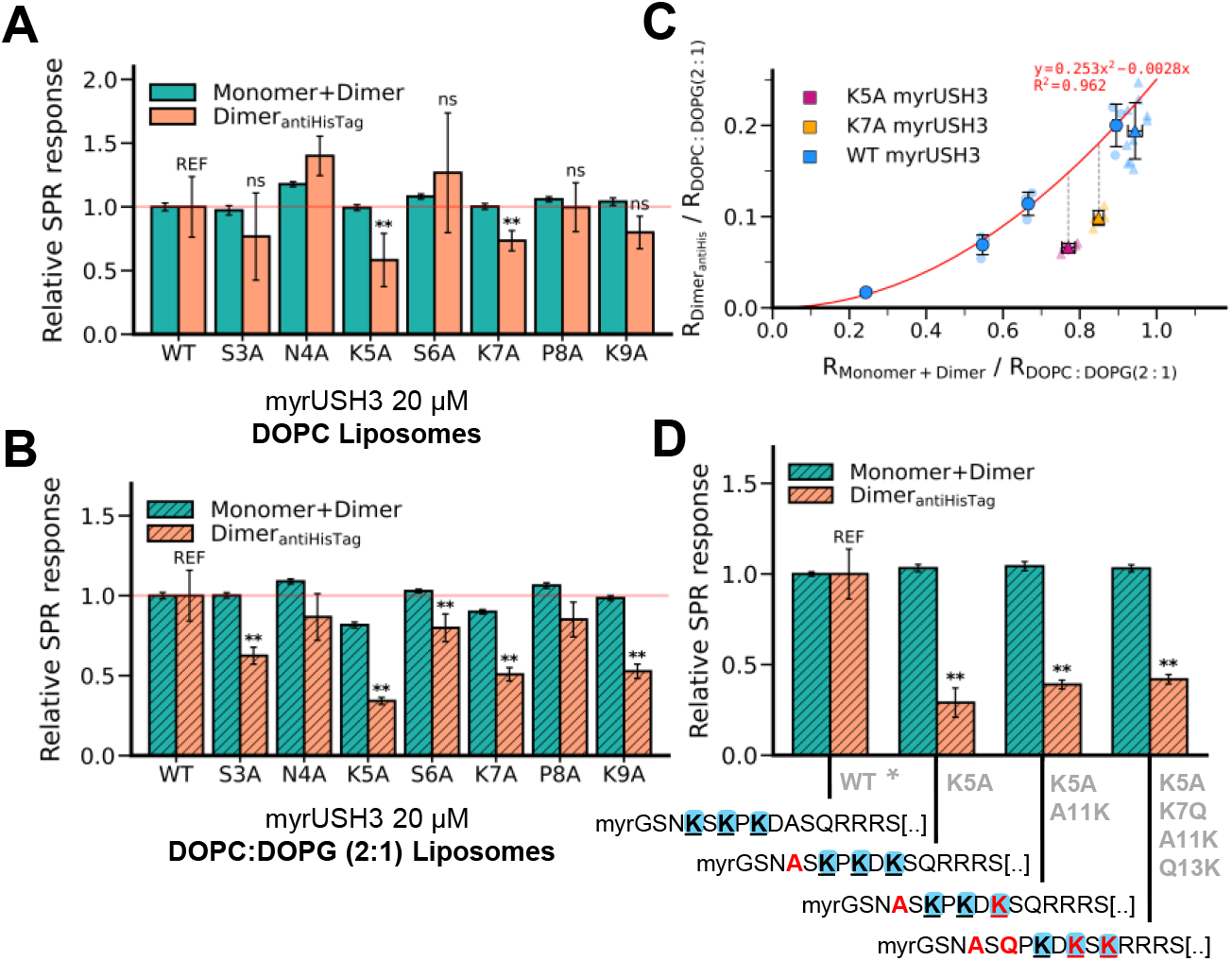
(A) and (B) Monomer+Dimer and Dimer_antiHisTag_ relative responses to WT for the X#A myrUSH3 mutants injected at 20 μM over (A) neutral DOPC liposomes and (B) negatively charged DOPC:DOPG (2:1) liposomes. Data expressed as mean ± SD, n = 4 (DOPC) n=5 (DOPC:DOPG (2:1)). (C) Dimer_antiHisTag_ relative response in terms of Monomer+Dimer relative response for K5A (magenta), K7A (orange) and WT myrUSH3 (blue). WT myrUSH3 was injected at a bulk concentration of 3, 6, 10 and 20 μM whereas the K#A mutants were injected at 20 μM. Data used originates from two different sensor chips indicated by the type of marker (circle or triangle). Data expressed as mean ± SD, n = 3 (circles) n ≥ 5 (triangles). (D) Monomer+Dimer and Dimer_antiHisTag_ relative responses to WT F64A myrUSH3 for the shifted K-cluster myrUSH3 F64A mutants injected at 20 μM over negatively charged DOPC:DOPG (2:1) liposomes. Data expressed as mean ± SD, n=6. Significant differences in the Dimer_antiHisTag_ with respect to WT construct are indicated with asterisks (Student’s t-test: *p < 0.1; **p < 0.05; ns: not significant; REF: reference).

### Monomer binding and dimerization are distinctly affected by K#A mutations

The fact that the mutation of the same residues hinders dimerization in charged and neutral lipids suggest that the effect does not depend on changes in the binding affinity of c-Src monomers. Reduction of the total charge by the K#A mutations lowers the affinity of the protein towards negatively charged lipids, and therefore reduces the density of bound monomers. In order to differentiate the effects of changes in protein density from variations in the intrinsic dimerization propensity of the individual K#A mutants we experimentally modified the steady-state protein density of WT myrUSH3 on the membrane by varying the bulk concentration at which it was injected between 3 μM and 20 μM (Fig. 2C). As expected for a dimer, the fraction detected by the antibody SPR response is proportional to the square of the intensity of the total protein density on the membrane, dominated by the reversibly bound monomer. Next, the K5A and K7A myrUSH3 mutants were injected at a fixed 20 μM bulk concentration. Both K#A mutants show a decreased steady-state response as compared to WT myrUSH3 but the population of dimer is reduced by ca. 40% for K7A and 50% for K5A taking as a reference the expected amount of dimer formed by the WT at the same protein density as K#A mutants. Thus, the observed decrease in dimer population cannot be explained only by a reduction on the protein density on the lipid surface. Interestingly, the steady-state response of the K5A and K7A mutants is not the same, despite having the same overall charge, suggesting that the position of the basic residues, and not just the overall charge, is important for primary binding and dimerization.

### Relative position of lysine residues with respect to the myristoylation site is important

Subsequently, we tested the effect of shifting the position of the lysine residue cluster (K-cluster). To this end we compared the dimerization on negatively charged lipids of myristoylated constructs in which residues K5 and K7 were mutated. We also introduced compensatory mutations downstream of the sequence, so that the total charge and residue composition were restored but the charged residues were increasingly displaced from the myristoylated N-terminus, while preserving the alternating pattern of positively charges (Fig. 2D). The mutants compared were: K5A, K5A+A11K and K5A+K7Q+A11K+Q13K. These proteins carried an additional mutation F64A, far from the SH4 domain, that resulted in increased stability of the mutants that, otherwise, tend to degrade spontaneously. As a control, we confirmed that the effect of the K5A mutation is similar in the WT and F64A mutants (cf. Fig. 2B and 2D). The compensatory A11K mutation restoring the total positive charge and thus the K-cluster, but maintaining the K5A mutation, does not recover the dimer formation of the WT form. Thus, confirming the important role of K5 beyond its contribution to the overall charge of the SH4 domain. A similar behavior is observed after the addition of mutation K7Q, together with the charge compensating mutation Q13K to the previous K5A/A11K. Thus, while mutation of individual lysine residues K5 and K7 cause a similar reduction in dimer formation, the effects are not additive. The importance of K5 and of its proximity to the myristoyl group in c-Src dimerization is further supported by the effect of the S3A mutation on myrUSH3 dimerization on charged lipids.

### Lysine to arginine mutations enhance c-Src dimerization

An isolated acidic residue, within or next to the lysine cluster, is found in position 10 of c-Src, Lyn, Blk and Frk, and in position 8 of Yes, Fyn and Yrk. We tested the importance of the single negatively charged residue D10 in c-Src by replacing it with alanine (D10A) and with a positively charged arginine (D10R) while maintaining the K-cluster intact (Fig. 3A-B). Arginine is known to interact with the phosphate and glycerol moieties in the lipid bilayer (21,22). The D10A and D10R c-Src mutants have a formal charge of +6 and +7 respectively, in the first 16 residues. The primary binding to negatively charged lipids is increased similarly in the D10R and D10A mutants, suggesting that the main effect is the elimination of the repulsion caused by the negatively charged residue. However, while the D10A has no effect on dimerization, the presence of the additional arginine residue causes a significant increase. The effect of the D10R mutation on binding to neutral lipids is much stronger than on negative lipids as dimerization is increased around 6-fold, while steady-state binding is similar to that of the D10A mutant.

**Figure 3.**
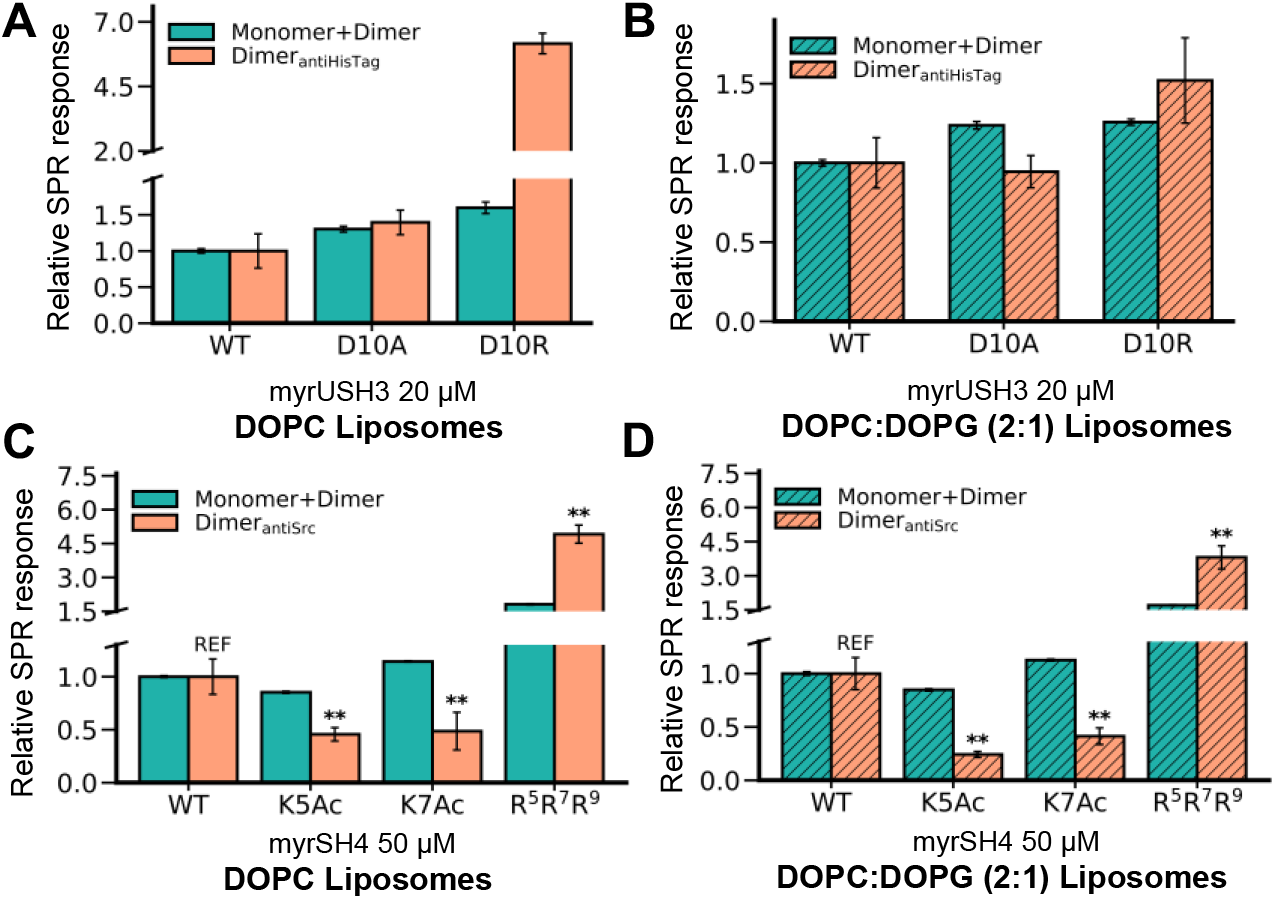
(A) and (B) Monomer+Dimer and Dimer_antiHisTag_ relative responses to WT for D10A and D10R myrUSH3 mutants injected at 20 μM over (A) neutral DOPC and (B) negatively charged DOPC:DOPG (2:1) liposomes. Data expressed as mean ± SD, n = 3 (DOPC) n=3 (DOPC:DOPG (2:1)). (C) and (D) Monomer+Dimer and Dimer_antiSrc_ relative responses to WT for acetylated (K5Ac and K7Ac) and triple arginine (R5R7R9) myrSH4 constructs injected at 50 μM over (C) neutral DOPC and (D) negatively charged DOPC:DOPG (2:1) liposomes. Data expressed as mean ± SD, n = 3 (DOPC) n=5 (DOPC:DOPG (2:1)). Significant differences in the Dimer_antiSrc_ of relative responses with respect to WT construct are indicated with asterisks (Student’s t-test: *p < 0.1; **p < 0.05; ns: not significant; REF: reference).

In order to further explore the role of the charged residues in the dimerization of myrUSH3, we used myristoylated synthetic peptides with residues 2-16 of c-Src, that form the entire SH4 domain (myrSH4) enabling the exploration of additional modifications, such as lysine acetylation, that eliminates the charge while preserving the rest of the side chain. In the absence of a His_6_Tag, we used a Src antibody recognizing its N-terminal region. As a control, we tested the effect of the K5A and S6A mutations individually and in combination (Fig. 4). The experiments with myristoylated peptides confirm that the K5A mutation inhibits persistent binding while S6A has no effect.

**Figure 4.**
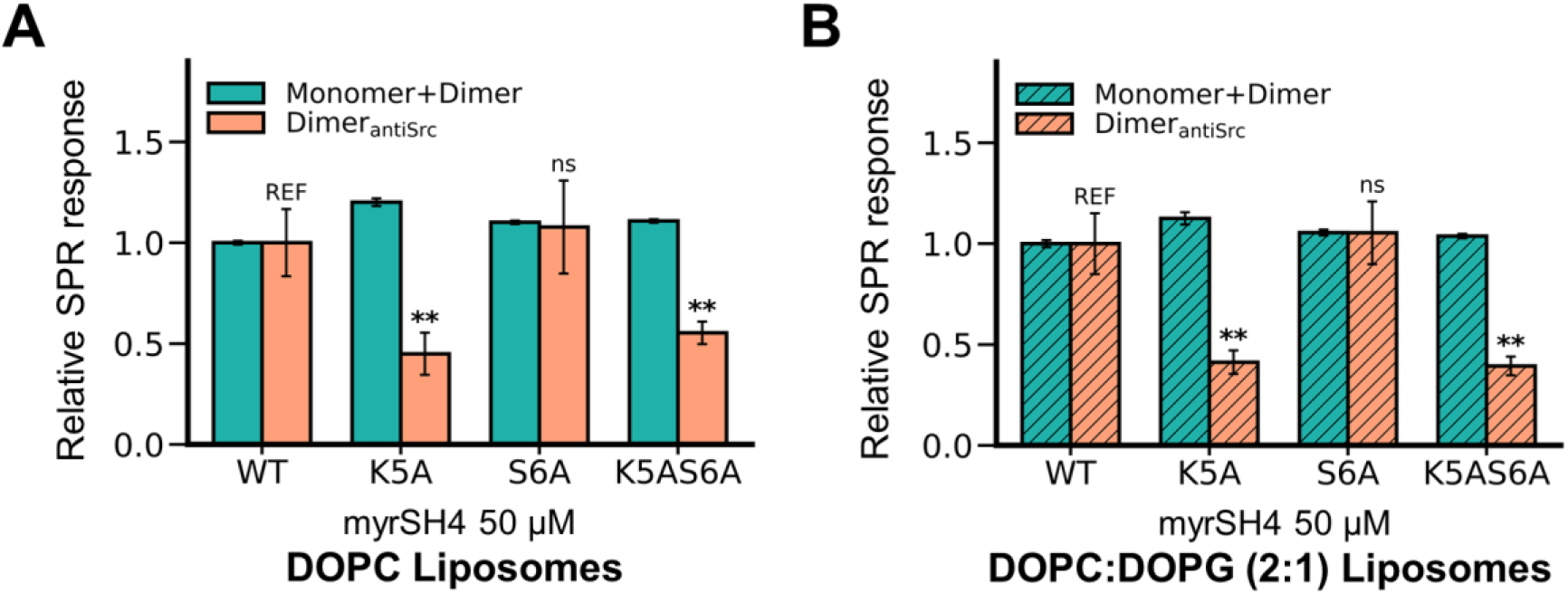
Monomer+Dimer and Dimer_antiSrc_ relative responses to WT for K5A, S6A and K5AS6A myrSH4 constructs injected at 50 μM over (A) neutral DOPC and (B) negatively charged DOPC:DOPG (2:1) liposomes. Data expressed as mean ± SD, n = 3 (DOPC) n=6 (DOPC:DOPG (2:1)). Significant differences in the Dimer_antiSrc_ relative responses with respect to WT myrSH4 are indicated by asterisks (Students t-test: *p < 0.1; **p < 0.05; ns: not significant, REF: reference).

We assessed the effect of acetylation of residues K5 and K7, that eliminate the positive charge, as well as the replacement of the three lysine residues by arginine to generate the R5R7R9 (3R) mutant that retains the same formal charge of wild type myrSH4 (Fig. 3C-D). It is reported that arginine maintains a stronger interaction with the lipid phosphate moieties at the membrane interface compared to lysine (21). Acetylation of either K5 or K7 significantly reduce the population of persistently bound peptide. On the contrary, a large increase is observed when lysine residues are replaced by arginine. The effects on zwitterionic or negatively charged lipids are very similar. The results with synthetic myristoylated peptides are consistent with those obtained with the entire myristoylated SNRE, in particular, the enhancement of dimer formation by arginine residues, as in the D10R mutant.

### Dimer formation in full length c-Src increases tyrosine phosphorylation in cells

We tested the effect of the 3R mutation in the context of full-length c-Src. To test in cell dimerization, wild type human c-Src and the 3R mutants were fused with FLAG and Myc tags at the C-terminus as described previously (6), and the various combinations of mutated or wild type proteins with the two tags were co-expressed in HEK293T cells. After cell lysis, proteins containing the FLAG tag were immunoprecipitated and tested for the co-precipitation of Myc-tagged c-Src (Fig. 5A). The 3R mutant shows a 50% increase in homodimer formation as compared to WT c-Src.

**Figure 5.**
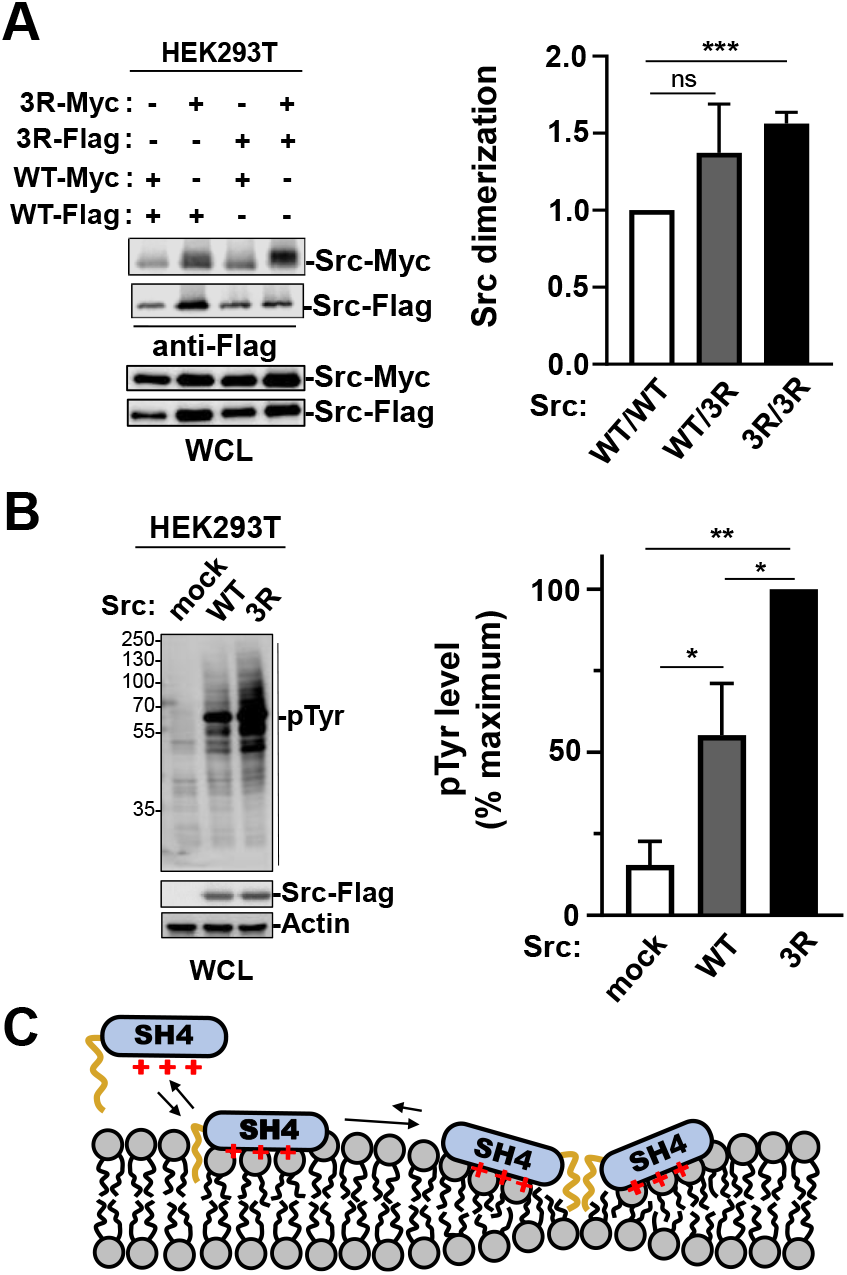
Mutations of lysine residues in positions 5, 7 and 9 to arginine (3R) results in enhanced dimer formation and enhanced tyrosine phosphorylation (A) Western blot of immunoprecipitated FLAG-tagged c-Src with anti-Myc and anti-FLAG antibodies. WCL is whole cell lysate. (B) Total tyrosine phosphorylation detected using an anti-pY antibody. The concentration of the two c-Src variants is very similar, as detected by anti-FLAG antibodies. Quantification is shown as the mean ± SEM; n=3; ns: p>0.05; *p<0.05; **p<0.01; Student’s t test. (C) Proposed model for a dimerization mechanism based on the distortion of the bilayer induced by the accumulation of positive charges together with the myristoyl group. This model would predict the formation of large clusters, which is observed in short peptides. Restriction of the oligomerization to dimers seems to be a consequence of interactions involving the adjacent Unique domain (not shown in the cartoon).

Since c-Src activation is known to involve trans autophosphorylation we predicted that increased dimerization should enhance the activation of c-Src. Consistently, the total tyrosine phosphorylation was increased by 100% in the 3R mutant as compared with wild type c-Src (Fig. 5B).

## Discussion

The synergistic effects of myristoyl insertion in the lipid bilayer and electrostatic interaction with the negatively charged lipids that mostly populate the inner leaflet of the cytoplasmic membrane have been demonstrated by the seminal work of McLaughlin and Resh (8,23–25). However, in this work, lipid-induced dimerization was not considered and electrostatic interactions and myristoyl insertion were treated as independent interactions collectively providing the binding energy to ensure stable binding although each of the individual interactions is not strong enough.

The present results suggest a model in which the entire myristoylated SH4 domain acts as a functional unit for lipid-mediated c-Src dimerization, in which the position of basic residues is at least as important as its number. Self-association driven by the positively charged SH4 domain is counterintuitive, as it should entail strong electrostatic repulsion. A high density of positive charges in the SH4 domain is shared with cell penetrating peptides. The interaction of HIV-Tat peptide with zwitterionic lipids has been simulated using molecular dynamics (26) showing that lysine and arginine side chains sequester neighbor lipid phosphate groups. When the density of cationic peptides increases, regions crowded with Tat peptides and phosphates are formed, as well as regions depleted from charged groups. The bilayer becomes thinner and the interaction with phosphate groups in the opposite lipid layer increases.

We suggest a similar model (Fig. 5C) may be occurring when the SH4 domain interacts with phospholipids, enhanced by the insertion of the myristoyl group in the lipid bilayer, leading to local changes in the width of the bilayer. Membrane deformations can propagate over several nanometers along the bilayer (27,28). At high protein densities, individual protein-induced deformations are no longer independent and result in membrane-mediated interactions, that can be attractive, repulsive, or non-monotonic. In particular, minimization of the protein-induced bilayer deformation drives protein lateral association (29). It should be noticed that membrane-mediated protein interactions are distinct from protein clustering driven by lipid-lipid phase separation. In our experiments, myrUSH3 dimerization is also observed in pure DOPC or in DOPC:DOPG mixtures that do not lead to lipid phase separation under the experimental conditions of our experiments.

Although the case of peripheral membrane proteins has been less studied, membrane-mediated association of amphipathic α-helical peptides has been predicted (28). The alternating lysine pattern in the SH4 domain is compatible with an amphipathic extended chain with the charged side chains pointing to the membrane phosphate groups. Suggestively, lateral interaction of the extended chains on the surface of the membrane may enable direct hydrogen-bonding between neighbor strands in a β-sheet-like arrangement. Lipid bound β-sheets have been observed in model cationic peptides with alternating lysine and not charged residues bound to anionic lipid bilayers (30). However, NMR chemical shifts of DMPC-bound myrSH4 indicates a random coil conformation for the lipid-bound peptide (31), although the presence of a minor population of self-associated peptide with broad signals may not be detectable.

The described model would predict extensive oligomerization/clustering of myristoylated c-Src. This is indeed observed in short peptides or constructs lacking the Unique domain. Our experiments using constructs containing the initial 85 residues of c-Src, i.e. the entire intrinsically disordered region, show that self-association stops at the dimer level. We speculate that steric interference between the disordered domains of c-Src molecules brought to very short distances through their N-terminus would have a strong cost (probably entropic) that hinders oligomerization beyond the dimer level.

Early work by Resh et al. had suggested the existence of a c-Src membrane receptor that could never be identified (32). The suggestion was based on the observation that membrane bound ^35^S-methionine labeled c-Src retained in pellets following ultracentrifugation could be competed off the lipid-bound fraction by an excess of myristoylated peptides containing only the initial 12 residues. These experiments were interpreted as evidence for competitive binding to a yet unidentified membrane receptor. However, on the light of our results, these experiments probably reflected the formation of persistently membrane bound c-Src dimers, which can be released by competition with the formation of heterodimers with myristoylated c-Src peptides. Indeed, using single molecule fluorescence photobleaching we had previously shown that doubly labeled homodimers were converted to single labeled heterodimers by incubating with unlabeled c-Src constructs (13).

Although dimeric c-Src represents only a minor species, c-Src activation is known to take place through trans-autophosphorylation (11). Therefore, modulating the formation of c-Src dimers may be a powerful regulatory mechanism. Known processes modifying the charge of the N-terminal lysine cluster of c-Src include their acetylation by CREB binding protein, described to result in the release of c-Src from the cell membrane (33). Similar alternating lysine clusters are observed in other SFKs (25): c-Src ^5^**K**S**K**P**K**D^10^, Lyn ^5^**K**S**K**G**K**D^10^, Yes ^5^**K**S**K**EN**K**^10^, Fyn ^7^**K**D**K**EAT**K**^13^. Except c-Src, the other SFK are palmitoylated, as well as myristoylated, so that positively charged groups to enhance primarily lipid anchoring are not required, suggesting that the role of the conserved cluster is to enable the association with other membrane anchored molecules of the same kind (e.g. SFK homodimers) or different (e.g. SFK mixed dimers). In the case of c-Src, the strong lipid binding of dimers, together with the intrinsic low membrane affinity of monomers, offer an efficient search mechanism to find the relevant membrane regions where signaling partners are located: weak lipid binding offers an efficient 2D diffusion mechanism, crossing corralling fences, while dimerization, induced by local signals (activated receptors, specific lipids…) can anchor and activate c-Src triggering its downstream signaling (12,13,18).

In the ensemble of the 58 human receptor tyrosine kinases, the juxtamembrane region protruding into the cytoplasm is enriched in positively charged residues. The initial 20 residues following the transmembrane helix contain 25% of basic residues and only 11% acidic ones (18). A positive cluster, containing lysine and arginine residues (^51^YA**RKR**NGQM^59^) is located in the small cytoplasmic tail extending from the small integral membrane protein SMIM30 encoded by lncRNA LINC0998. SMIM30 induces c-Src/Yes membrane anchoring promoting hepatocellular carcinoma (34). Although the binding mode has not been demonstrated, it is known to involve the N-terminal region of c-Src and, on the light of the above results, we suggest that it involves lipid-mediated hetero-dimerization/oligomerization involving SMIM30 C-terminal tail and one or more Src molecules through their SH4 domain.

Interestingly a similar positive cluster motif is present in K-Ras isoform 2B forming nanoclusters (35). The membrane anchoring hypervariable disordered region is located at the C-terminus, and like c-Src, depends on the insertion of a hydrophobic chain (farnesyl, instead of myristoyl) and the electrostatic interaction of a cluster of positively charged residues with negatively charged lipids. The farnesylated cysteine is preceded by the sequence ^180^**K**S**K**T**K**^184^.

While the abundance of positively charged residues in the juxtamembrane region of integral membrane proteins has been long recognized, our results suggest that, as in the case of c-Src, clusters of basic residues may be a general motif involved in membrane-mediated protein-protein interactions.

## Materials and Methods

### Myristoylated c-Src variants expression and purification

Myristoylated c-Src constructs were obtained by co-expression of the N-myristoyl transferase (NMT) enzyme along the c-Src USH3-His6Tag construct in a pETDuet-1 (Novagen) plasmid. The mutations were introduced using QuickChange II XL Site Directed Mutagenesis Kit (Agilent) and Q5 High-Fidelity PCR Kit (New England BioLabs). Plasmids were transformed in *E. coli* Rosetta™ (DE3)pLysS (Novagen). Bacteria cells were grown in LB medium supplemented with chloramphenicol (25 µg/mL) and ampicillin (100 µg/mL) at 37ºC until an OD600nm of ∼0.6. A freshly prepared solution of myristic and palmitic acid (Sigma) (200 µM final concentration/each) and fatty acid free BSA (Sigma) (600 µM final concentration) was added to the cell culture. The lipid-BSA solution was prepared by adding one equivalent of NaOH, heating at 65ºC until complete lipid dissolution and adjusting the final pH to 8. Protein expression was induced with 1mM of IPTG (Nzytech) and performed for 5 h at 28ºC. 1h after induction start, 6 g/L of glucose was added to the cell culture. Cells were harvested at 4000 rpm for 20 min. Bacteria was lysed in buffer containing 20 mM Tris·HCl, 300 mM NaCl, 10 mM Imidazole, pH 8, 1 % Triton X100 (Sigma), 1x Protein Inhibitor Cocktail (Sigma), 1 mM PMSF (Sigma), 25μg/mL lysozyme (Sigma) and 5μg/mL DNAseI (Roche). Cells were sonicated on ice and centrifuged at 25000 rpm for 45 min. Ni-NTA affinity chromatography was performed using a 1mL Ni-NTA cartridge (GE Healthcare). The protein was eluted with buffer containing 20 mM Tris·HCl, 300 mM NaCl, 10 mM Imidazole, 400 mM imidazole, 0.02 % Triton X100 (Sigma) at pH 8. The final purification step consisted of a size exclusion chromatography in a Superdex 75 26/60 (GE Healthcare) using phosphate buffer (50 mM NaH_2_PO_4_/Na_2_HPO_4_, 150 mM NaCl, 0.2 mM EDTA, pH 7.5). The protein was concentrated with a Vivaspin 20, 5 kDa MWCO concentrator (Sigma Aldrich). The purity of the protein was established by HPLC in a BioSuite pPhenyl 1000RPC 2.0 × 75 mm; 10 μm column coupled to mass spectrometry (ESI-MS), confirming the complete protein myristoylation (Table 1). All proteins were flash-frozen and stored at −80ºC. Myristoylated SH4 domain peptides (myrSH4) were synthesized by SynPeptide Co., Ltd (Shangai, China).

**Table 1.**
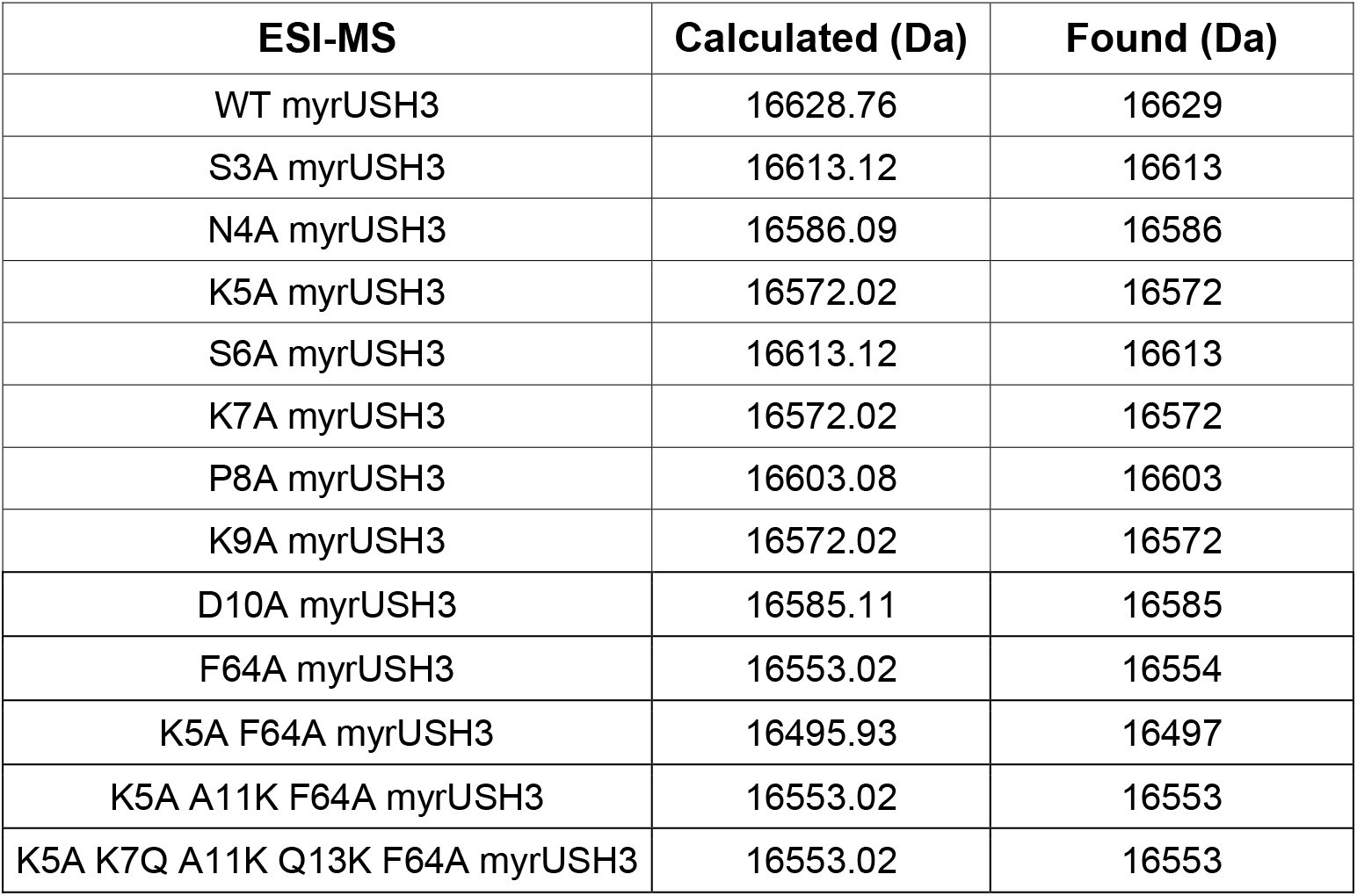
Results from the mass spectrometry analysis (ESI-MS) of the different myrUSH3 constructs.

### Preparation of liposomes

1,2-dioleoyl-sn-glycero-3-phosphocoline (DOPC) (TebuBio) and 1,2-dioleoyl-sn-glycero-3-phospho(1’-rac-glycerol) (sodium salt) (DOPG) (Sigma) were dissolved in chloroform. Lipid compositions used were DOPC and DOPC:DOPG (2:1) at 4mM. The organic solvent was evaporated under a nitrogen stream. Lipid films were rehydrated with vortexing in phosphate buffer 50 mM NaH_2_PO_4_/Na_2_HPO_4_, 150 mM NaCl, 0.2 mM EDTA, pH 7.5 (SPR assays). Large unilamellar vesicles were prepared by extrusion using a Mini-Extruder (Avanti Polar Lipids). The lipid suspension was extruded 15 times through a 100 nm polycarbonate filter. The mean diameter of the liposomes was verified by Dynamic Light Scattering (Zetasizer Nanoseries S. Malvern instruments). Liposomes were used within two days to avoid lipid oxidation.

### Surface Plasmon Resonance assays

All measurements were carried out in a Biacore T200 instrument (GE Healthcare) at 25ºC. A 2D-carboxylmethyldextran sensor chip (Xantec) was used. All the flow cells, except for the reference, were modified by a covalent attachment of phytosphingosine (TebuBio) to allow the capture of liposomes. An amine-coupling procedure was performed with 1 mM of phytosphingosine in freshly prepared 20 mM sodium acetate buffer at pH 6.7. The running buffer for all experiments consisted of 50 mM NaH_2_PO_4_/Na_2_HPO_4_, 150 mM NaCl, 0.2 mM EDTA, pH 7.5. DOPC and DOPC:DOPG (2:1) liposomes at 1 mM concentration were coated over the phytosphingosine containing flow cells by a 20 s injection at 10 µL/min. In case the distinct flow cells contained different types of lipids, the DOPC liposomes were injected in the first available flow cell to avoid anionic lipid migration towards the neutral liposomes through the flow cells. The reference cell and possible uncovered surface in the liposomes flow cells were blocked with 1 mg/ml of BSA at 50 µL/min for 20 s. Mass transport effects were minimized by injecting the myristoylated c-Src variants at 50 µL/min. Protein concentration ranged from 3 µM to 20 µM for the protein density experiments and it was fixed to 20 µM for the mutants analysis. The myrSH4 peptides were injected at 50 µM. The protein was allowed to associate for 60 s while the dissociation lasted 350 s. The antibody (antiHis_6_Tag or antiSrc antibodies (both SantaCruz Biotechnology)) was injected at 1:5 dilution in running buffer for 60 s at 30 µL/min. n≥3 experiments were performed for the c-Src variants, each time in randomized order. The surface was regenerated with two pulses (30 s at 100 µL/min) of Isopropanol:50 mM NaOH (2:3) solution followed by a 20 mM CHAPS (Sigma) or 40mM Octyl-beta-Glucoside (Sigma) pulse. Each cycle started with freshly captured liposomes. All data was double referenced (reference channel and baseline subtraction (0μM concentration)) and analyzed using Biacore T200 3.0 Evaluation software (GE Healthcare). Experiments were performed in two sensor chips, reproducible as indicated in Fig. 2C.

### Data and statistical analysis

The different mutants in Fig. 2 and 3 are compared relatively to WT myrUSH3 or myrSH4 as indicated per equations 1 and 2.

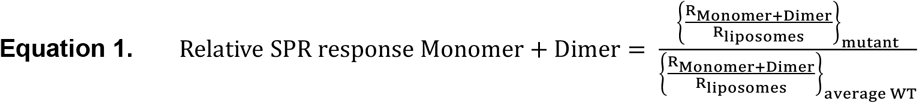

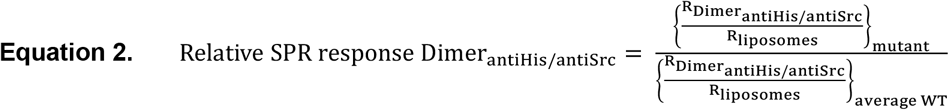

Where R is the SPR response observed in the corresponding sensorgram. All the statistical analysis were performed using the Python scientific library SciPy (SciPy: Open-Source Scientific Tools for Python, http://www.scipy.org, last accessed Apr 12, 2022). If not otherwise stated, the analysis was performed against the WT c-Src construct. Unpaired two-sample t-test (two-sided) was conducted.

### Src dimerization in HEK293T cells

PcDNA3 vectors expressing human Src-myc and Src-Flag were described in (Aponte et al. (6)). pcDNA3-Src K5R/K7R/K9R (Src 3R) mutants were obtained by PCR using the QuickChange Site-Directed Mutagenesis Kit (Stratagene); forward 5’:CCCTTCACCATGGGTAGCAACAGGAGCAGGCCCAGGGATGCCAGCCAGCGGCGCCGC, reverse 5’:GCGGCGCCGCTGGCTGGCATCCCTGGGCCTGCTCCTGTTGCTACCCATGGTGAAGGG. HEK293T cells (ATCC, Rockville, MD) were cultured and transfected with indicated Src constructs for 24h as described (Aponte et al. (6)). Immunoprecipitation and immunoblotting of Src proteins were performed as described in (Aponte et al. (6)) by using anti-Flag coupled beads (Pierce, #A36797) for Src precipitation and anti-Flag (M2, Sigma) and anti-Myc (9B11, Ozyme) antibodies for Src immunoblotting. Cellular protein tyrosine phosphorylation was performed by immunoblotting of total cell lysate (20 μg) using anti-pTyr antibody (4G10, a gift form Prof. Mangeat; CRBM) and anti-Actin (Cell Signaling Technology). Signal quantification and analyses were performed using ImageJ and GraphPad Prism respectively. Data are presented as the mean ± SEM from 3-4 independent experiments (two-tailed t-test; p values: ns p>0.05, *p≤0.05, **p≤0.01, ***p≤0.001).

## Acknowledgments

We acknowledge extensive discussions with Miguel Arbesú and the help of Roger Martínez in the preparation of mutants. This work has been funded by the Spanish Agencia Estatal de Investigación (projects BIO2016-78006R, co-funded with Regional Development Funds, PID2019-104914RB-I00, and PDC2021-121629-I00, financed by European Union Next Generation Funds), Fundació Roviralta, Ministry of Science and Innovation (project PID2019-104914RB-I00), la Ligue Nationale Contre le Cancer (LNCC), Montpellier SIRIC Grant «INCa-DGOS-Inserm 6045», and the Agènce Nationale de Recherche (FUZZY-SRC ANR-21-CE13-0011).

## References

1. Thomas SM, Brugge JS. Cellular functions regulated by SRC family kinases. Vol. 13, Annual Review of Cell and Developmental Biology. 1997. p. 513–609.

2. Yeatman TJ. A renaissance for SRC. Vol. 4, Nature Reviews Cancer. 2004. p. 470–80.

3. Allgayer H, Boyd DD, Heiss MM, Abdalla EK, Curley SA, Gallick GE. Activation of src kinase in primary colorectal carcinoma: An indicator of poor clinical prognosis. Cancer. 2002;94(2):344–51.

4. Arbesú M, Maffei M, Cordeiro TN, Teixeira JMC, Pérez Y, Bernadó P, et al. The Unique Domain Forms a Fuzzy Intramolecular Complex in Src Family Kinases. Structure. 2017;25(4):630-640.e4.

5. Le Roux AL, Mohammad IL, Mateos B, Arbesú M, Gairí M, Khan FA, et al. A Myristoyl-Binding Site in the SH3 Domain Modulates c-Src Membrane Anchoring. iScience. 2019;12:194–203.

6. Aponte E, Lafitte M, Sirvent A, Simon V, Barbery M, Fourgous E, et al. Regulation of Src tumor activity by its N-terminal intrinsically disordered region. Oncogene. 2022;41(7):960–70.

7. Kamps MP, Buss JE, Sefton BM. Rous sarcoma virus transforming protein lacking myristic acid phosphorylates known polypeptide substrates without inducing transformation. Cell. 1986;45(1):105–12.

8. McLaughlin S, Aderem A. The myristoyl-electrostatic switch: a modulator of reversible protein-membrane interactions. Vol. 20, Trends in Biochemical Sciences. 1995. p. 272–6.

9. McLaughlin S. The electrostatic properties of membranes. Vol. 18, Annual review of biophysics and biophysical chemistry. 1989. p. 113–36.

10. Ben-Tal N, Honig B, Peitzsch RM, Denisov G, McLaughlin S. Binding of small basic peptides to membranes containing acidic lipids: Theoretical models and experimental results. Biophys J. 1996;71(2):561–75.

11. Barker SC, Kassel DB, Weigl D, Huang X, Luther MA, Knight WB. Characterization of pp60c-src Tyrosine Kinase Activities Using a Continuous Assay: Autoactivation of the Enzyme Is an Intermolecular Autophosphorylation Process. Biochemistry. 1995;34(45):14843–51.

12. Le Roux AL, Busquets MA, Sagués F, Pons M. Kinetics characterization of c-Src binding to lipid membranes: Switching from labile to persistent binding. Colloids Surfaces B Biointerfaces. 2016;138:17–25.

13. Le Roux AL, Castro B, Garbacik ET, Garcia Parajo MF, Pons M. Single molecule fluorescence reveals dimerization of myristoylated Src N-terminal region on supported lipid bilayers. ChemistrySelect. 2016;1(4):642–7.

14. Irtegun S, Wood RJ, Ormsby AR, Mulhern TD, Hatters DM. Tyrosine 416 Is Phosphorylated in the Closed, Repressed Conformation of c-Src. PLoS One. 2013;8(7):e71035.

15. Spassov DS, Ruiz-Saenz A, Piple A, Moasser MM. A Dimerization Function in the Intrinsically Disordered N-Terminal Region of Src. Cell Rep. 2018;25(2):449-463.e4.

16. Maffei M, Arbesú M, Le Roux AL, Amata I, Roche S, Pons M. The SH3 domain acts as a scaffold for the N-terminal intrinsically disordered regions of c-Src. Structure. 2015;23(5):893–902.

17. Ahler E, Register AC, Chakraborty S, Fang L, Dieter EM, Sitko KA, et al. A Combined Approach Reveals a Regulatory Mechanism Coupling Src’s Kinase Activity, Localization, and Phosphotransferase-Independent Functions. Mol Cell. 2019;74(2):393-408.e20.

18. Pons M. Basic Residue Clusters in Intrinsically Disordered Regions of Peripheral Membrane Proteins: Modulating 2D Diffusion on Cell Membranes. Physchem. 2021;1(2):152–62.

19. Dwivedi M, Mejuch T, Waldmann H, Winter R. Lateral Organization of Host Heterogeneous Raft-like Membranes Altered by the Myristoyl Modification of Tyrosine Kinase c-Src. Angew Chemie -Int Ed. 2017;56(35):10511–5.

20. Himeno H, Shimokawa N, Komura S, Andelman D, Hamada T, Takagi M. Charge-induced phase separation in lipid membranes. Soft Matter. 2014;10(40):7959–67.

21. Li L, Vorobyov I, Allen TW. The different interactions of lysine and arginine side chains with lipid membranes. J Phys Chem B. 2013;117(40):11906–20.

22. Wu Z, Cui Q, Yethiraj A. Why do arginine and lysine organize lipids differently? Insights from coarse-grained and atomistic simulations. J Phys Chem B. 2013;117(40).

23. Silverman L, Resh MD. Lysine residues form an integral component of a novel NH2-terminal membrane targeting motif for myristylated pp60(v-src). J Cell Biol. 1992;119(2):415–26.

24. Murray D, Hermida-Matsumoto L, Buser CA, Tsang J, Sigal CT, Ben-Tal N, et al. Electrostatics and the membrane association of Src: Theory and experiment. Biochemistry. 1998;37(8):2145–59.

25. Silverman L, Sudol M, Resh MD. Members of the src family of nonreceptor tyrosine kinases share a common mechanism for membrane binding. Cell Growth Differ. 1993;4(6):475–82.

26. Herce HD, Garcia AE. Molecular dynamics simulations suggest a mechanism for translocation of the HIV-1 TAT peptide across lipid membranes. Proc Natl Acad Sci U S A. 2007;104(52):20805–10.

27. Andersen OS, Koeppe RE. Bilayer thickness and membrane protein function: An energetic perspective. Vol. 36, Annual Review of Biophysics and Biomolecular Structure. 2007. p. 107–30.

28. Kondrashov O V., Galimzyanov TR, Jiménez-Munguía I, Batishchev O V., Akimov SA. Membrane-mediated interaction of amphipathic peptides can be described by a one-dimensional approach. Phys Rev E [Internet]. 2019 Feb 1;99(2):022401. Available from: https://link.aps.org/doi/10.1103/PhysRevE.99.022401

29. Kondrashov O V., Galimzyanov TR, Pavlov K V., Kotova EA, Antonenko YN, Akimov SA. Membrane Elastic Deformations Modulate Gramicidin A Transbilayer Dimerization and Lateral Clustering. Biophys J. 2018;115(3):478–93.

30. Hädicke A, Blume A. Binding of cationic model peptides (KX) 4 K to anionic lipid bilayers: Lipid headgroup size influences secondary structure of bound peptides. Biochim Biophys Acta -Biomembr. 2017;1859(3):415–24.

31. Scheidt HA, Klingler J, Huster D, Keller S. Structural Thermodynamics of myr-Src(2-19) Binding to Phospholipid Membranes. Biophys J. 2015;109(3):586–94.

32. Sigal CT, Zhou W, Buser CA, McLaughlin S, Resh MD. Amino-terminal basic residues of Src mediate membrane binding through electrostatic interaction with acidic phospholipids. Proc Natl Acad Sci U S A. 1994;91(25):12253–7.

33. Huang C, Zhang Z, Chen L, Lee HW, Ayrapetov MK, Zhao TC, et al. Acetylation within the N- and C-terminal domains of src regulates distinct roles of STAT3-mediated tumorigenesis. Cancer Res. 2018;78(11):2825–38.

34. Pang Y, Liu Z, Han H, Wang B, Li W, Mao C, et al. Peptide SMIM30 promotes HCC development by inducing SRC/YES1 membrane anchoring and MAPK pathway activation. J Hepatol. 2020;73(5):1155–69.

35. Barceló C, Paco N, Beckett AJ, Alvarez-Moya B, Garrido E, Gelabert M, et al. Oncogenic K-ras segregates at spatially distinct plasma membrane signaling platforms according to its phosphorylation status. J Cell Sci. 2013;126(20):4553–9.

